# Volitional control of a recurrent brain computer interface via covert reinforcement of endogenous reward networks

**DOI:** 10.64898/2026.10.01.755922

**Authors:** Jacob G. McPherson, Robert R. Miller, Eberhard E. Fetz, Steve I. Perlmutter

**Affiliations:** Program in Physical Therapy, Washington University in St. Louis, St. Louis, MO; Dept. of Anesthesiology, Washington University in St. Louis, St. Louis, MO; Program in Neurosciences, Washington University in St. Louis, St. Louis, MO; Dept. of Biomedical Enginering, Washington University in St. Louis, St. Louis, MO; Dept. of Neurobiology and Biophysics, University of Washington, Seattle, WA

## Abstract

Recurrent brain–computer interfaces (rBCIs) deliver targeted stimulation to one brain region based on neural activity recorded and processed in real-time at another. In a volitionally controlled rBCI, the user must also learn to intentionally produce the triggering neural activity, a process typically facilitated by an external reward. However, reliance on an external reward introduces a delay between the triggering event and reinforcement that can slow association between the two and, ultimately, control of the rBCI (particularly for neural activity not directly related to an observable behavior such as a goal-directed movement). The constraint to an external reward also limits studies of neural dynamics during natural, free behavior. In contrast, delivering activity-dependent stimulation directly to the brain’s own reward centers—i.e., covert reinforcement—would bypass the need for external feedback, enabling precise temporal coupling between triggering events and reward and allowing for unrestrained rBCI use. Here, we show that rats can gain volitional control of an rBCI using activity-dependent electrical stimulation of the medial forebrain bundle as a sole reinforcer. Animals successfully learned to modulate motor unit discharge rates as well as specific electrocorticographic frequency bands (beta and gamma) to self-trigger stimulation. Furthermore, rats demonstrated flexible control by independently modulating distinct spectral features from the same cortical site. These findings suggest that covert reinforcement of endogenous reward networks can drive operant conditioning of specific neural signals during unrestrained behavior, opening new avenues for adaptive BCIs and closed-loop psychiatric therapies.

## Introduction

The realization that animals could gain direct, volitional control of neural activity through operant conditioning ^1^ revolutionized study of the brain. Now borne out across multiple species, networks, and scales, the brain’s ability to directly control its own synaptic interactions through operant conditioning has become central both to interrogating basic mechanisms of neural plasticity and to developing new treatments for neurological injuries and diseases through brain computer interfaces ^2–4^.

Recurrent brain computer interfaces (rBCIs) deliver activity-dependent electrical stimulation to the brain ^5^. Neural activity is recorded from one brain region and processed in real-time using programmable decision criteria to trigger stimulation of a different region of the brain. Such closed loop systems can bridge lost neural connections or promote reorganization of circuits made maladaptive or weakened by injury or disease ^6–13^. A largely unexplored use of rBCIs is to deliver activity-dependent stimulation to limbic or modulatory centers of the brain to ‘covertly’ ^14, 15^ produce behavioral adaptations and/or to reinforce learning and memory ^16–20^. Such systems would be ideally suited to empower two applications for which BCIs hold considerable, yet unrealized, potential: mechanistic studies of plasticity during free behavior and translational studies focusing on treatment of psychiatric illnesses (i.e., cognitive, affective, and mood disorders)^21^.

In all rBCIs, the brain must learn both to integrate the function(s) of the artificial connections with those of existent neural pathways and to use the reorganized circuitry. In a volitionally controlled rBCI, there is also an explicit need for the brain to learn to control the neural activity that triggers stimulation – a process typically facilitated by operant conditioning. However, operant conditioning for BCI applications requires a constrained environment in which the reinforcing feedback (e.g., juice, food pellet, cursor movement) can be rapidly delivered upon detection of the operant behavior. This constraint often prevents study of neural dynamics during the free behaviors for which most BCIs are intended. An rBCI that delivers reinforcing activity-dependent stimulation to the brain’s endogenous reward circuitry to condition a desired operant behavior – covert reinforcement – could potentially overcome this constraint by obviating the need for external reinforcement. It would also enable more precise synchronization between detection of the operant behavior and delivery of reinforcement, facilitating the association between the two.

Electrical stimulation of limbic structures also holds promise for managing otherwise refractory psychiatric illnesses ^17, 22–27^. To date, however, electrical stimulation of these structures for psychiatric applications has overwhelmingly been delivered in an open-loop manner. That is, stimulation is delivered independently of the presence or absence of symptoms or particular brain states. Such uncontrolled stimulation may have complications, including unintended effects when symptoms are not present, habituation, and/or excessive stimulation over time.

A more optimal approach would be an autonomous, closed-loop system that implements an unconscious homeostatic modulation of affective neural networks. And indeed, if a signal from a particular brain region could predict the onset of symptoms of an affective disorder, an rBCI could recognize this aberrant activity and automatically begin stimulation to alleviate the symptoms. Unfortunately, however, appropriate signals have yet to be identified. Thus, a first step may be to gain volitional control of a particular neural signal to enable activity-dependent self-stimulation of endogenous reward circuitry.

Here, we sought proof-of-principle for whether an rBCI delivering electrical stimulation to the brain’s endogenous reward circuitry could reinforce a desired operant behavior, enabling the rBCI user to gain volitional control of the triggering activity. We first demonstrate that rats can learn to control the frequency of motor unit action potential discharge within a narrow temporal window to receive activity-dependent electrical stimulation of the medial forebrain bundle (MFB), a large, evolutionarily conserved fiber pathway that interconnects limbic, hypothalamic, and other brain regions and produces a strong reinforcing stimulus ^28^. Next, we demonstrate that rats can learn to volitionally modulate cortical population activity (power in a specific local field potential frequency band) to receive reinforcing stimulation of the MFB. Finally, we demonstrate that a rat can independently modulate power in different local field potential frequency bands recorded at the same cortical site to receive MFB stimulation.

## Methods

Experiments were performed in 2 adult female Long Evans rats (280 – 350 g). All procedures were approved by the Institutional Animal Care and Use Committee of the University of Washington.

### Surgical Procedures

Both rats underwent an identical surgical procedure to implant a deep brain stimulating electrode, epidural recording electrodes, and EMG electrodes. Surgical procedures were performed using sterile techniques.

Rats were anesthetized with 2-3% isoflurane in oxygen and given preoperative ketoprofen, enrofloxacin, dexamethasone and local anesthetic. Physiological vital signs were monitored continuously during the procedure, and temperature was maintained at 37 ℃. Once a deep, surgical plane of anesthesia was achieved, the scalp was incised rostrocaudally and reflected. Between 6 and 8 small burr holes were drilled unilaterally in the skull over sensorimotor cortex exposing the dura mater. Multi-stranded stainless steel wires (30 μm diameter; Cooner Wire, Chatsworth CA, USA) were inserted through the holes and glued in place epidurally to record electrocorticographic (ECoG) signals (**Fig. 1, inset**). A single custom-made tungsten stimulating electrode (50 μm diameter) was then implanted through an additional burr hole to target the medial forebrain bundle (MFB) contralateral to the ECoG electrodes (**Fig. 1, inset**). Coordinates for the MFB were 2.8 mm caudal, 1.7 mm lateral and 7.8 mm ventral to bregma ^29^.

**Figure 1.**
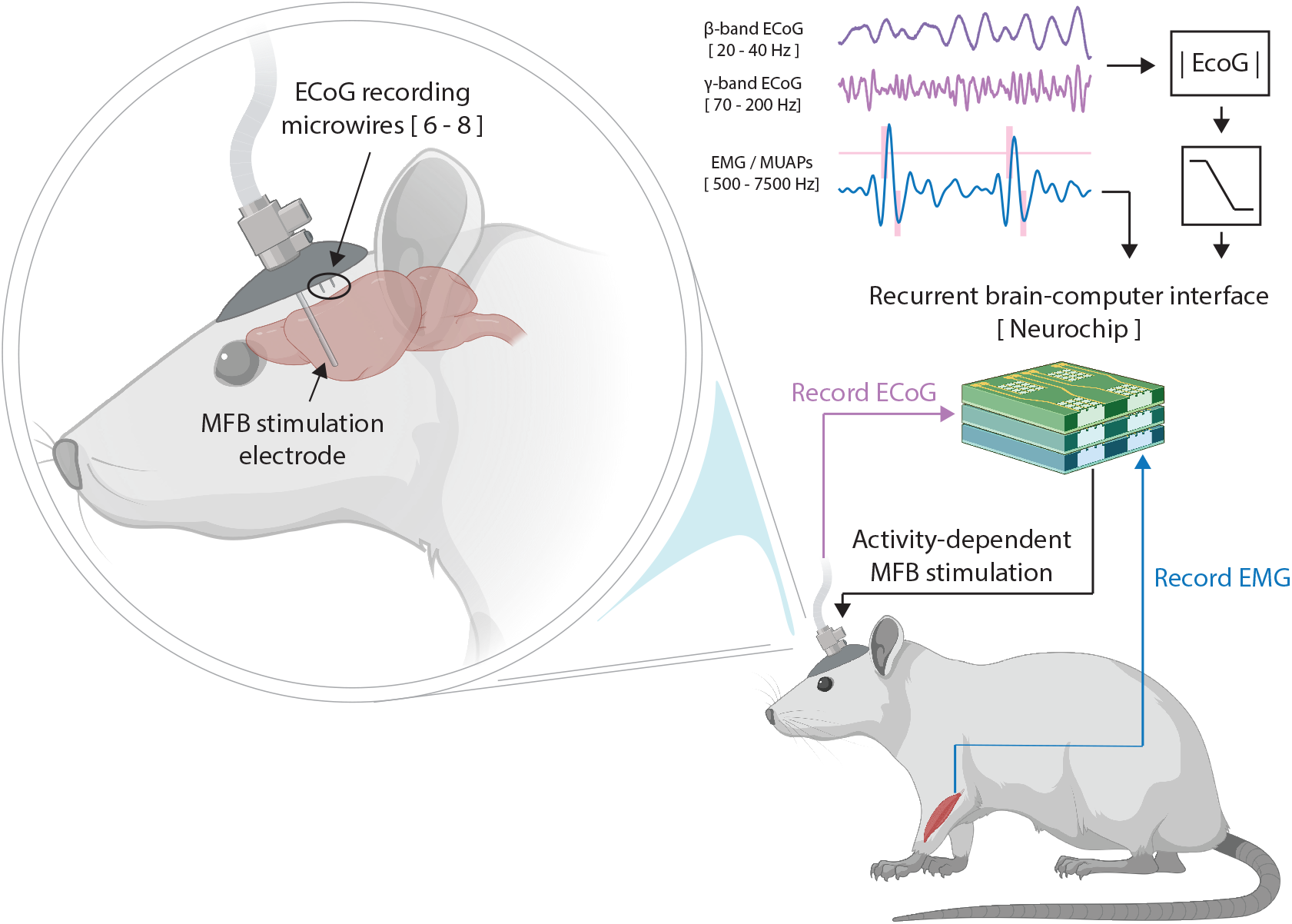
Schematic overview of the rBCI. *Left*: enlarged depiction of the head-fixed components of the rBCI, consisting of a deep brain stimulation electrode in the MFB, 6-8 epidural ECoG wires, and a skull-mounted headcap with electrical connector. The wire extending from the headcap was connected via flexible cable and commutator to the Neurochip, located outside of the behavioral testing arena (not shown). *Right*: animals separately modulated the power in specific ECoG bands (β and γ bands) and EMG, recorded via epidural cortical electrodes and intramuscular forelimb extensor electrodes, respectively, to receive activity-dependent electrical stimulation of the MFB. The Neurochip brain-computer interface was used to process neural activity and EMG in real-time and deliver MFB stimulation upon detection of qualifying events.

Wire leads from the ECoG and MFB electrodes were routed to a custom connector that was affixed to the skull via bone screws and dental acrylic (**Fig. 1, inset**). Electrical grounding wires were wrapped around screws on the left and right sides of the skull. In the same procedure, 2 pairs of bipolar, multistranded, stainless steel wires (Cooner Wire, Chatsworth CA, USA) were implanted into forelimb extensor muscles contralateral to the ECoG electrodes (**Fig. 1**). The wires were then routed subcutaneously to the same head-mounted connector containing the ECoG and MFB leads. Buprenorphine was given postoperatively as an analgesic, and animals were allowed at least 1 week of recovery before experiments began.

### Electrophysiology and operant conditioning paradigms

All physiological signals were recorded with the Neurochip, which is a miniature, autonomous, battery-powered rBCI ^30–32^. The Neurochip was also used to deliver stimulation through the MFB electrode. A schematic representation of the setup is shown in **Fig. 1**. ECoG signals were filtered for the beta frequency band (β = 20-40 Hz) and/or the gamma frequency band (γ = 70–200 Hz), rectified, and smoothed in real-time over 100 ms sliding windows. Both the β and γ bands are prominent signals correlated with the activity of neurons near the recording electrodes. EMG signals were filtered from 500 – 7500 Hz, enabling detection of single motor unit action potentials (MUAPs) via a real-time dual-window time-amplitude discriminator.

For both animals, the experimental sequence was as follows: (1) sessions of MFB stimulation triggered by a lever press; (2) sessions of operant conditioning of forelimb extensor EMG to trigger MFB stimulation; and (3) sessions of operant conditioning of ECoG to trigger MFB stimulation. An explicit number of experimental sessions was not defined *a priori* for each phase, both because the learning rate for such tasks is unestablished and because we predicted that it would vary between animals. Sessions ranged from 1-2 h each and no more than 1 session was conducted per day. All sessions were divided into 10 min epochs, alternating between reinforcement periods in which MFB stimulation was delivered and extinction periods in which no stimulation was delivered (i.e., “catch trial” periods). An audible tone cued the animal to the reinforcement periods. MFB stimulation was delivered as a single burst of stimuli upon each trigger event (10 biphasic pulses, 150 uA/phase, 167 Hz), followed by a 2 s rate-limiting refractory period to prevent excessive stimulation. All experimental sessions occurred in 10” x 12” arenas in which the animals could move freely.

The initial lever press sessions served two primary functions: to introduce the animals to the presence and sensation of MFB stimulation and to verify the reinforcing efficacy of the MFB electrode site itself. Proper placement of the electrode was confirmed when the animals consistently pressed the lever to receive MFB stimulation without a food reward ^33, 34^. Following the lever press sessions, animals were transitioned to the covert reinforcement sessions. EMG from a forelimb extensor muscle was used as the initial physiological trigger for MFB stimulation. In this paradigm, the Neurochip detected discrete motor unit action potentials and delivered a burst of MFB stimulation when 6-8 events occurred within a 100 ms window (**Fig. 2a**). Once an animal successfully established volitional control of MFB stimulation using EMG as a trigger, the paradigm was transitioned to operantly condition increases in ECoG activity. In this case, MFB stimulation was delivered when the rectified, smoothed ECoG signal exceeded a pre-set threshold voltage, typically ∼2 standard deviations from the baseline signal amplitude (e.g., **Fig. 3a**).

**Figure 2.**
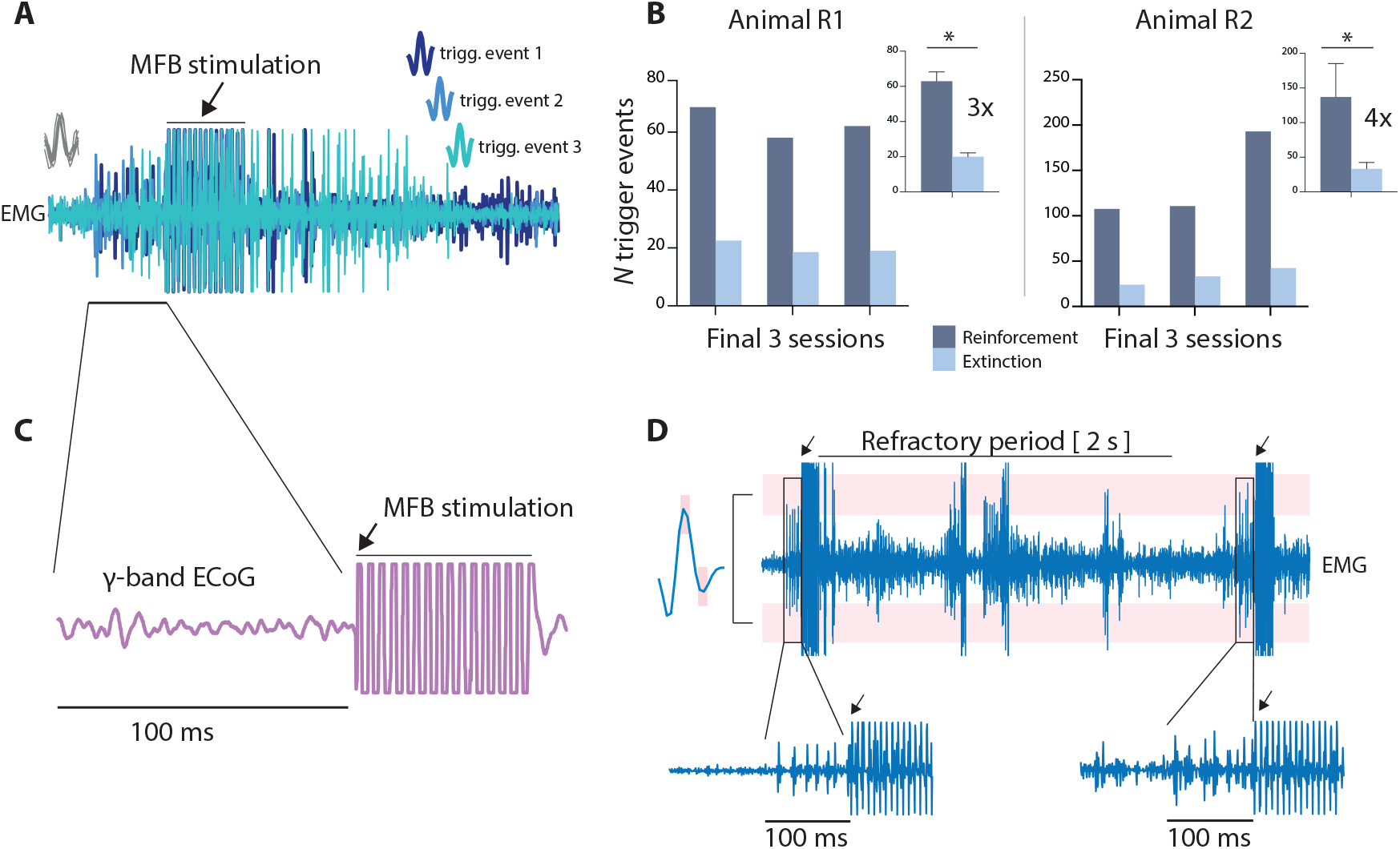
Motor unit-based control of the rBCI. *A*: time-aligned overlay of three representative EMG records containing motor unit activity that triggered MFB stimulation. *B*: number of motor unit-based triggering events per animal during each of the last three conditioning sessions. Epochs when MFB stimulation was available are termed “reinforcement” (gray) and epochs when MFB stimulation was unavailable are termed “extinction” (blue). *B, insets*: average number of motor unit-based triggering events per animal over the last three reinforcement and extinction epochs. Both animals produced significantly more qualifying events during reinforcement epochs (animal R1: 63±5 vs. 20±2, *p* = 0.001; animal R2: 137±48 vs. 33±9, *p* = 0.02). *C*: ECoG activity recorded during production of a motor unit-based stimulation trigger, highlighting the lack of correlation between the two signals. *D*: MFB stimulation does not itself produce EMG activity, which could lead to spurious triggering events. *Left*: trace of a single motor unit action potential satisfying the dual time-amplitude window criteria. *Top, main*: representative record of EMG activity (3 s) containing two triggering events and two contingent MFB stimulation events. Pink overlays depict amplitude thresholds through which motor unit action potentials were required to pass as one criterion for stimulation. Black arrows indicate onset of MFB stimulation. Following each MFB stimulation event, a refractory period of 2 s was imposed, indicated by the horizontal line. Following each stimulation event, EMG activity fell outside of the qualifying range. Potentially qualifying events during the refractory period are ignored. Note that the second of the two qualifying trigger events appears at a delay relative to the end of the refractory period, further indicating that the MFB stimulus did not trigger rebound EMG activity capable of satisfying the triggering criteria. *Bottom*: zoomed depiction of 2 motor unit-based triggering events.

**Figure 3.**
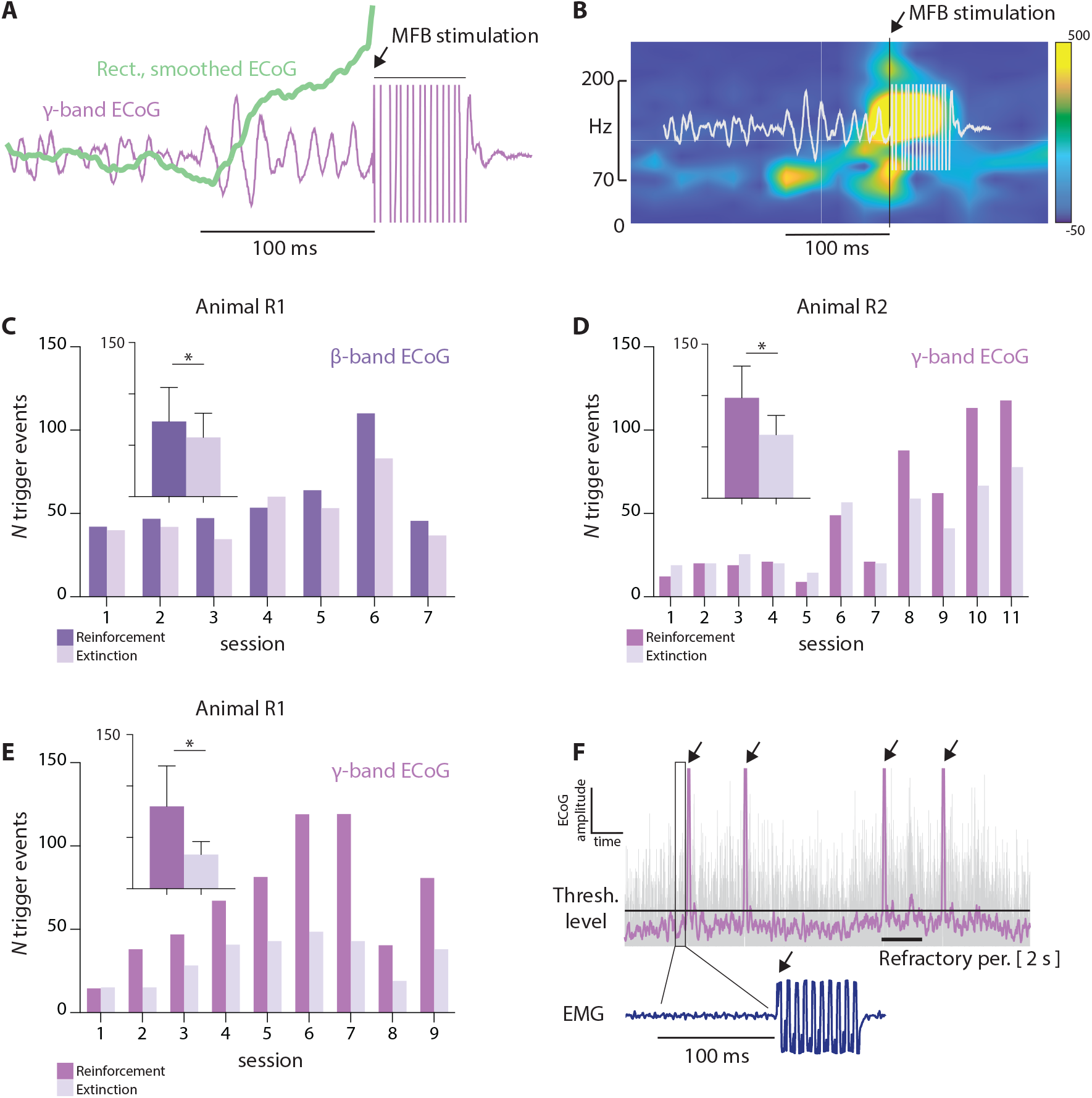
ECoG-based control of the rBCI. *A*: representative ECoG-based MFB stimulation trigger event. Purple: raw γ-band ECoG signal; green overlay: rectified, smoothed γ-band ECoG, used to identify qualifying triggering events. Increase in γ-band activity over 100 ms precedes onset of MFB stimulation, indicated by black arrow. *B*: time-frequency representation of broadband ECoG prior to a qualifying γ-band trigger event, highlighting that the triggering activity is confined spectrally to the conditioned band (here, γ-band, 70-200 Hz) and temporally to 100 ms immediately prior to stimulation. *C*: animal R1 began the ECoG-based rBCI experiments by learning to modulate power in the β-band. The main plot depicts the number of qualifying β-band triggering events across the first 7 sessions, separated by reinforcement epochs (MFB stimulation available; dark) and extinction epochs (MFB stimulation unavailable; light), 10 min each. The inset shows the average number of qualifying trigger events over the last 3 reinforcement and extinction sessions, respectively (73±33 vs. 58±23; *p* = 0.05). *D*: same as *C*, but for animal R2 and γ-band ECoG sessions. The average number of qualifying trigger events over the last 3 reinforcement and extinction sessions, respectively, was 98±31 vs. 62±19 (*p* = 0.02). *E*: ECoG- triggered MFB stimulation for animal R1 when switched from β-band control to γ-band control at the same cortical recording site. The plot layout is the same as *C, D*. Over the last three sessions, the average number of qualifying γ-band triggers during reinforcement epochs (80±39) significantly exceeded the number produced during extinction epochs (33±13) (*p* = 0.05). *F*: ECoG-triggered MFB stimulation does not produce recursive triggers after cessation of the refractory period. Purple: rectified, smoothed γ-band ECoG; gray: raw γ-band ECoG; horizontal line: threshold level for triggering stimulation; arrows: onset time of stimulation. Note that filtered ECoG decreases below threshold prior to each subsequent trigger. Bottom: representative EMG trace depicting the absence of correlated activity during the triggering period.

### Statistical analyses

The primary outcome measure for both EMG and ECoG-triggered stimulation was the number of trigger events that occurred during reinforcement periods vs. extinction periods. Potential differences in the occurrence of trigger events were assessed separately for each animal using unpaired, two-way t-tests. Differences were considered statistically significant when *p-*values were ≤ 0.05. Given the exploratory, proof-of-principle nature of this study, a power analysis was not performed *a priori* to determine sample size.

Experimentalists were not blinded to whether an animal was in a reinforcement or extinction period at any given point in time. However, the neither the EMG nor ECoG operant conditioning sessions required intervention from the experimentalist. Thus, no confounds were introduced by the lack of blinding. Data analysis was performed by an investigator not directly involved in data collection. Numerical and graphical data in text and figures are presented as mean ± standard deviation.

## Results

### EMG control of MFB stimulation

We first used food rewards to train the animals to press a lever. Upon each press, the experimentalist delivered stimulation through the MFB electrode. Once the lever task was learned – indicating that the animals associated the task with reward – we discontinued the use of food incentives and only delivered MFB stimulation in response to each lever press, similar to the foundational demonstrations of reinforcing deep brain stimulation in rats ^34^. Both animals consistently pressed the lever in the absence of food rewards, indicating that the stimulation was indeed interpreted as reinforcing and thereby also confirming appropriate electrode placement.

We then asked whether animals could gain direct, volitional control of MFB stimulation. Because activation of the triceps muscle was an implicit requirement of the previously acquired lever press task, we facilitated the learning process by using triceps EMG activity as the initial trigger for stimulation. Unlike the lever press task, however, we required the animals to modulate specific features of the EMG activity to receive reinforcing stimulation (the lever press task imposed no constraints on EMG activity or the movement strategy adopted by each animal). Specifically, animals were required to modulate motor unit discharge rate. **Figure 2a** depicts an overlay of 3 representative motor unit-based trigger events.

Over time, both animals learned to volitionally control motor unit activity to receive MFB stimulation. Animal R1 gained proficiency in the task—indicated by consistently more qualifying triggers during reinforcement epochs than extinction epochs—within ∼8 conditioning sessions, and for animal R2, ∼9 sessions. For both animals, we collected 3 additional motor unit sessions after volitional control initially appeared to be established. These sessions served two primary purposes.

First, they were intended to provide additional confirmation that volitional control of stimulation had been achieved and was not merely a chance occurrence of the animals generating more trigger events during reinforcing epochs than during extinction epochs. The raw number of trigger events per epoch during the 3 sessions following acquisition of volitional control is shown for each animal in **Fig. 2b**. Summary data from these sessions are depicted in **Fig. 2b, insets**. Across these sessions, animal R1 generated ∼3X more trigger events per reinforcement epoch than per extinction epoch (63±5 vs. 23±2; *p* = 0.001; **Fig. 2b**), while animal R2 produced ∼4X more trigger events during reinforcement epochs than during extinction epochs (137±48 vs. 33±9; *p* = 0.02; **Fig 2b**). These findings support the conclusion that volitional control was established.

The second purpose of these 3 additional sessions was to determine whether the reinforcing efficacy of MFB stimulation decreased over time. However, we found no evidence of habituation, reward satiation, or mental fatigue: the number of trigger events during reinforcing epochs did not decrease (and increased over the last 3 sessions in animal 2) and the number of trigger events during extinction epochs did not increase.

Given that our goal was to train animals to gain volitional control over ECoG activity to trigger stimulation, we sought to verify that production of motor unit-based stimulation triggers was not associated with ECoG activity satisfying the triggering criteria. That is, it was important to understand whether control of the two trigger sources was independent. If not, then it could be argued that the animals merely perpetuated the motor unit-triggered stimulation approach instead of learning to volitionally control ECoG through MFB-stimulation driven operant reinforcement. To assess this possible confound, we computed averages of ECoG aligned to each EMG-based stimulation trigger (**Fig. 2c**). The lack of increased ECoG in the 100 ms immediately prior to each stimulus during the EMG conditioning sessions confirmed the independence of the two signals when EMG was used as the triggering source.

Finally, although EMG-triggered volitional control appeared to be robust, it was possible that MFB stimulation itself could have evoked the triggering EMG activity, giving the appearance of volitional control. Such an effect was unlikely, both because the stimulation site is not a component of descending motor pathways and because the refractory period should have prevented ‘rebound’ stimulation even if it evoked EMG activity satisfying the triggering criteria. And indeed, if stimulation-evoked EMG triggered rebound MFB stimulation, we would predict it to occur at the nominal rate-limited 1-2 Hz. Nevertheless, to confirm that MFB stimulation did not directly cause EMG activity resulting in a rebound stimulus, we verified that EMG activity decreased below triggering threshold prior to the subsequent trigger event and stimulus. A representative example is show in **Fig. 2d**.

### ECoG control of MFB stimulation

After verifying that both animals could gain volitional control of motor-unit activity to trigger MFB stimulation, we asked whether the animals could gain volitional control of ECoG activity to receive MFB stimulation. In animal R1, we began by operantly conditioning β-band ECoG; in animal R2, γ-band. **Fig. 3a** depicts a representative example of a stimulus burst triggered by γ-band ECoG. Band-limited raw data are shown in purple, with the rectified, smoothed data used for triggering stimulation overlaid in green. **Fig 3b** depicts a time-frequency representation of the same triggering event, highlighting the increases in γ-band power leading to stimulation.

Over time, both animals gained volitional control of ECoG to receive MFB stimulation. Somewhat surprisingly, acquisition of this skill required a comparable number of conditioning sessions as when the operant was motor unit activity. Qualitatively, animal R1 gained volitional control of MFB stimulation via β-band modulation within ∼5-7 conditioning sessions, as evidenced by the greater number of qualifying triggers during reinforcement than extinction epochs over this span. Animal R2 appeared to gain volitional control of MFB stimulation via γ-band ECoG after ∼8 conditioning sessions (**Fig. 3c, d**).

As with the motor unit paradigm, we conducted additional experimental sessions for both animals after volitional control initially appeared to be established. Summary data from these sessions are depicted in the insets of **Fig. 3c, d**. Across these sessions, animal R1 generated 1.3X more β-band trigger events per reinforcement epoch than per extinction epoch, a modest but significant increase (73±33 vs. 58±23; *p* = 0.05; **Fig. 3c**), while animal R2 produced, on average, 1.6X more γ-band trigger events during reinforcement epochs than during extinction epochs (98±31 vs. 62±19; *p* = 0.02; **Fig 3d**).

We then sought additional confirmation that MFB stimulation was under direct, volitional control via purposeful modulation of ECoG activity. First, we reasoned that one distinguishing feature of volitional control would be the ability to flexibly adapt to a new cortically derived operant. To that end, we terminated β-band control of MFB stimulation in animal R1 and instead required control of γ-band ECoG to receive stimulation. Remarkably, the animal began to consistently produce more qualifying triggers during reinforcement than extinction epochs in only its second session—earlier than its performance in both the motor unit and β-band paradigms—and averaged >2X more trigger events during reinforcement than extinction epochs over its last 3 conditioning sessions (80±39 vs. 33±13, *p* = 0.05; **Fig. 3e**). Its control of the rBCI via γ-band ECoG was also considerably more robust than with β-band control, as evidenced by the greater ratio of reinforcement to extinction period triggers across all sessions (2.0±0.55 for γ vs. 1.2±0.16 for β; *p*=001). Next, we verified that MFB stimulation itself did not lead to persistently elevated ECoG activity sufficient to trigger involuntary reinforcement loops. A representative example is depicted in **Fig. 3f** (top), showing subthreshold ECoG activity prior to a series of sequential trigger events, even when one event closely follows conclusion of the prior refractory period.

Although the EMG-triggered average ECoG (**Fig. 2c**) suggested that motor unit and cortical control were independent, we questioned whether the animals might have perpetuated use of movement-associated triggers during ECoG sessions instead of directly producing qualifying ECoG activity. Thus, we created ECoG-triggered averages of EMG activity, aligned by stimulation onset. As is evident in the representative example of **Fig. 3f** (bottom), EMG activity was stable during the 100 ms window immediately prior to MFB stimulation, despite the concurrent increase in ECoG amplitude required to trigger the stimulation event. Video recordings of motor unit and ECoG sessions also revealed distinct behavioral patterns consistent with independence of the motor unit and ECoG-based trigger sources. Specifically, after the animals acquired volitional control of the rBCI via ECoG, they remained largely stationary in the testing arenas during reinforcement and extinction epochs (also reflected in the negligible pre-stimulation EMG activity). In contrast, the animals were highly active during the motor unit sessions, as would be expected to maximize the number of reinforcing stimuli delivered in that paradigm.

We then turned to the question of habituation, reward satiation, and mental fatigue. The additional conditioning paradigm completed by animal R1, combined with its faster learning rate in the γ-band task, provided the opportunity to study the potential impact of prolonged stimulation on reward efficacy. We investigated the potential interaction of stimulation duration/frequency and reward efficacy at two timescales: within a day and across multiple days. On two days (separated by a week) we conducted two full rBCI sessions on the same day (sessions 3 and 4, and sessions 6 and 7 in **Fig. 3e**). On both occasions, the average number of reinforcement period triggers during the second session of the day was greater than or equal to that of the first session, suggesting that reward efficacy did not decrease over short timescales.

To investigate potential changes in reward efficacy occurring over longer timescales, we considered the 7 rBCI sessions [3,9] in animal R1—i.e., those occurring after it first generated more stimulation triggers during reinforcement than extinction epochs via γ-band ECoG. We found that the number of rewarded trigger events progressively increased after apparent initial acquisition of volitional control (session 2), peaking during sessions 6 and 7 before declining during sessions 8 and 9 (**Fig. 3e**). It is difficult to infer whether this decrease was due to habituation, reward satiation, or mental fatigue, or whether it was related to other (albeit unknown) environmental or physiological factors. Regardless, however, the number of reinforcement period trigger events in each of these final two sessions remained ∼2X greater than the corresponding number of triggering events during extinction epochs, suggesting that control accuracy did not decrease.

## Discussion

Our primary findings are that (1) rats can learn to increase the frequency of motor unit action potential discharge to receive reinforcing stimulation of the MFB, (2) rats can learn to volitionally modulate power in specific ECoG frequency bands to receive reinforcing stimulation of the MFB, and (3) rats can independently modulate power in different ECoG frequency bands recorded at the same cortical site to receive reinforcing MFB stimulation. These findings indicate that focal stimulation of the MFB can mediate covert operant reinforcement in an rBCI task. Together, these findings have implications for advancing our understanding of neural plasticity during free behavior and for studying and potentially treating a wide range of psychiatric illnesses.

### Independence of motor unit vs. cortical control of the rBCI

In this proof of principle study we tested multiple rBCI control strategies to probe whether MFB stimulation could provide sufficient reinforcing efficacy to serve as a generalizable, application-agnostic approach to providing covert reinforcement ^14, 15^ for rBCI implementation. We predicted that the learning rate for motor unit-based rBCI control would be faster than for ECoG-based control. This prediction was based on recognition that the force production required for the initial lever press task also implicitly required the animals to increase their EMG activity, which would have been realized through increased motor unit recruitment and rate modulation. However, such a difference was not borne out by the empirical data. On the contrary, each rat’s initial ECoG-based learning rate was comparable to its respective motor unit-based rate.

It was possible that prior acquisition of motor unit-based control could have facilitated acquisition of volitional ECoG control, which might have otherwise been slower. Three lines of evidence argue against this possibility. First, we observed a decrement in performance when switching between spinal and cortical control strategies, indicative of the need to re-learn the rBCI task. Second, we found no evidence of β or γ-band modulation time-locked to the triggering motor unit activity. And third, we found no evidence of increased motor unit discharge rate time-locked to the triggering ECoG activity; in fact, animals reduced their gross activity level as they became more proficient with ECoG-based control. In contrast to the apparent independence of acquiring motor unit and cortical control of the rBCI, we speculate that acquiring a second mode of cortical control was facilitated by prior acquisition of cortical control at the same site.

### Motor unit vs. cortical control of the rBCI: control accuracy and generalizability

Although it is not possible to dissociate unintentional trigger events from intentional trigger events in this paradigm, motor unit-based control of the rBCI appeared to be more accurate than ECoG-based control. This conclusion is supported by the higher ratio of reinforcement period trigger events to extinction period trigger events during motor unit sessions compared to ECoG sessions, an effect driven by an increased number of extinction period trigger events during the ECoG sessions rather than a systematic increase in the number of reinforcement period trigger events. Presumably, the cortical population activity satisfying the triggering criteria more closely resembled ongoing, unrelated physiological activity than the triggering motor unit activity, leading to unintentional triggers during extinction epochs. Such an effect could have contributed to the increased number of γ-band ECoG triggers during reinforcement epochs for animal R1, as well.

### Multi-dimensional control of cortical population activity from a single site

It is understood that the brain can learn to independently modulate the firing rate of different neurons and to independently modulate different features of population activity accessed at distinct sites. It has also been demonstrated that the brain can simultaneously control multiple ensembles of 2-4 neurons accessed at nearby sites ^35^. However, the extent to which the brain can learn to control multiple features of neural activity at the same site is unestablished. The present study was not designed to probe this question directly. However, our finding that a rat could successfully, albeit not robustly, gain volitional control of γ-band ECoG power after acquiring control of β-band ECoG power at the same cortical site supports the notion that such multi-dimensional control may be possible. Future studies will be necessary to understand these actions, including whether the brain can exert control of two or more features simultaneously (e.g., increase power in one frequency band while reducing power in another) and/or the extent to which the brain can rapidly and accurately switch between control strategies.

### Neural plasticity during free behavior

The foundational demonstrations of the reinforcing potential of deep brain self-stimulation required rats to press a lever to receive stimulation ^33, 34^. Although the rats were physically unrestrained, this arrangement imposes numerous experimental constraints. There is now much interest in studying the adaptability of brain activity during truly free behavior, as it is expected that the richness and flexibility of neural dynamics are masked by environmentally restricted conditions. However, studies during free behavior require a means of delivering temporally precise reinforcement. It may be difficult to achieve this with lever presses in the cage. Here, we demonstrated that activity-dependent MFB stimulation can be delivered in real-time using an rBCI. We also established that its reinforcing efficacy is sufficient to rapidly condition multiple types of neural activity or behaviors. And although we utilized control variables accessed from movement related structures, this approach could be generalized to numerous other structures or networks for which studies during free behavior would address fundamental unresolved questions in the field.

### Activity-dependent stimulation of affective networks as a target for potential clinical translation

In a typical operant reinforcement paradigm, the desired operant behavior is not explicitly to receive the reinforcing feedback; reinforcement is a means to an end, not the end itself. However, from the perspective of a volitionally controlled rBCI for psychiatric applications, this reconceptualization of the canonical operant conditioning paradigm is useful. To that end, we chose to stimulate the MFB less for its potential therapeutic efficacy in the context of a specific psychiatric illness than to determine if it could promote covert reinforcement of cortical population activity in a volitionally controlled rBCI task. That being said, deep brain stimulation of the MFB is actively being investigated as a potential treatment for major depressive disorder ^28, 36–38^, obsessive compulsive disorder ^39^, and bipolar disorder ^40^.

Other stimulation sites within the limbic system are also being investigated as potential treatments for various psychiatric illnesses ^16, 21, 41^. Among these are the nucleus accumbens, the subgenual cingulate gyrus and ventral striatum, the anterior limb of the internal capsule, the ventral caudate, the lateral habenula, and the subthalamic nucleus. In principle, any of these sites could be targeted for stimulation in an rBCI, although presumably different sites will be more or less optimal for managing different conditions and multi-site stimulation may be necessary to achieve adequate symptom relief ^26^. It is also possible that stimulation of many of these (or other) sites could covertly reinforce the production of specific features of neural activity to gain volitional control of an rBCI. Aside from the present findings, however, only the nucleus accumbens has been investigated in this context ^19^.

Regardless of the stimulation site(s), electrophysiological identification of brain states potentially indicative of affective symptoms continues to be a major barrier to the use of rBCIs for psychiatric applications. Although research in this domain is increasing ^17, 24, 42–45^, it will likely remain necessary to trigger symptom alleviating stimulation from volitionally controlled neural activity for the foreseeable future (as opposed to homeostatic trigger sources). Motor unit action potentials and motor cortical population activity, as used here, presumably do not play a direct role in the pathophysiology of psychiatric illnesses. Therefore, we expect that volitional control of neural activity from different sites may be more appropriate for this application. However, given the relative ease with which animals and humans can gain control of these trigger sources ^3, 4, 46^, they may prove to be convenient for testing the reinforcing and/or therapeutic efficacy of new rBCI systems.

A final consideration for long-term use of a volitionally controlled rBCI delivering covert reinforcement is the potential for habituation, reward satiation, and/or mental fatigue ^33^. Here, such effects would manifest as the production of fewer stimulation triggering events over time, either within a session or across sessions. Although animal R1 did exhibit a decline in the number of trigger events over its last two sessions of ECoG-based control, it is not possible to unequivocally determine whether this finding was in fact due to a lower reinforcing efficacy of MFB stimulation or to other factors. Regardless, the rats in this study were self-stimulating at far higher frequencies (1 event/∼5 s) than would be expected in a clinically applied therapy, suggesting that a decrease in reinforcing efficacy could potentially be avoided with imposition of additional rate-limiting measures.

